# The vaginal microbiota shapes local and systemic inflammation during *Chlamydia trachomatis* infection in non-human primate

**DOI:** 10.1101/2025.02.20.638673

**Authors:** Cindy Adapen, Louis Réot, Natalia Nunez, Claude Cannou, Romain Marlin, Julien Lemaitre, Sabrine Lakoum, Ségolène Diry, Léo d’Agata, Wesley Gros, Anne-Sophie Gallouët, Marco Leonec, Laetitia Bossevot, Nathalie Dereuddre-Bosquet, Roger Le Grand, Marie-Thérèse Nugeyre, Elisabeth Menu

**Affiliations:** Université Paris-Saclay, Inserm, CEA; “Immune diseases, microbiology and innovative therapies” (IDMIT/UMRS1184); Fontenay-aux-Roses & Le Kremlin-Bicêtre, France; Life&Soft, 92260 Fontenay-aux-Roses, France; MISTIC group, department of Virology, Institut Pasteur, Paris, France

**Keywords:** Vaginal microbiota, *Lactobacillus*, *Chlamydia trachomatis* infection, inflammation, neutrophils

## Abstract

Chlamydiosis, a sexually transmitted infection induced by *Chlamydia trachomatis* (CT), increases local inflammation (cytokine production, recruitment of neutrophils). Vaginal microbiota (VM) is involved in the regulation of local immune responses and STI acquisition where *Lactobacillus spp* are associated with protection. In this study, the VM of cynomolgus macaques was enriched with *Lactobacillus crispatus* after local metronidazole treatment followed by repeated intravaginal inoculations of CT. Metronidazole treatment induced drastic VM modifications (decrease abundance of several anaerobes and increase of natural *Lactobacillus spp)*. After CT exposure, a non-persisting CT infection and no association between *L. crispatus* enrichment and a lower CT susceptibility were detected. However, serum specific anti-CT IgG production was higher in treated animals. High concentration of blood cytokines after CT infection was observed in untreated animals, whereas treated animals exhibited an increased production of cervicovaginal cytokines. Blood neutrophils were more mature and activated after CT infection in both groups. These results suggest an efficient local immune response associated with a controlled systemic inflammation (low cytokine production and higher anti-CT IgG titer) upon CT infection in *L. crispatus* treated animals. This study confirmed that VM composition can be a powerful tool to modulate local immune responses to STIs.

**Authors summary:** *Chlamydia trachomatis* (CT) is a bacterial pathogen responsible for a highly prevalent sexually transmitted infection (STI), particularly among young women. Often asymptomatic, untreated infections can ascend to the upper genital tract, causing inflammation and, in severe cases, infertility. Environmental factors within the female reproductive tract, including the vaginal microbiota, can influence susceptibility to infection by modulating local immune responses. A non-*Lactobacillus* spp.-dominated microbiota has been associated with an increased risk of CT acquisition. Using the relevant *in vivo* model of cynomolgus macaques, which closely resemble humans in terms of immune responses and female reproductive tract anatomy, we investigated the impact of vaginal *Lactobacillus crispatus* colonization on CT infection. The presence of *L. crispatus* did not prevent CT acquisition but was associated with increased CT-specific antibody levels. Moreover, systemic inflammation was higher in the non-*Lactobacillus*-treated group, whereas the opposite trend was observed in the cervicovaginal compartment. This study highlights the critical role of microbiota composition in shaping host immune responses to pathogen infection.

## Introduction

Sexually transmitted infections (STIs) continue to represent a major public health issue. In many countries, 15 to 24 years old sexually active women have the highest risk of *Chlamydiosis*. *Chlamydia trachomatis* (CT) is an obligate intracellular gram-negative bacterium and serovar D to K are responsible for genital infection. Around 70% to 90% of CT infection in women are asymptomatic and without active screening, individuals stay infected and continue to spread the pathogen [1]. Asymptomatic or symptomatic CT infection can induce chronic inflammation in the lower genital tract, but the infection can also progress to the upper tract causing, in severe cases, sterility[1,2].

CT preferentially infects the endocervical columnar epithelial cells of the female reproductive tract (FRT). Upon infection, pro-inflammatory cytokines increase in the FRT, resulting in a local inflammation[3–7]. A Th1 response with IFNγ production is necessary to clear CT infection whereas anti-CT IgG/IgA are associated with a reduced cervical CT burden[8]. Polymorphonuclear leukocytes (PMN) like other innate immune cells are recruited to the FRT during CT infection[4]. Neutrophil accumulation and prolonged survival have been linked to immunopathology and are not directly necessary for bacterial clearance[9,10].Moreover, an accumulation of activated vaginal neutrophils (CD62L^low^CD16^high^) with a prolonged lifespan was observed in women with a non-*Lactobacillus* dominant microbiota linking vaginal microbiota composition and neutrophil phenotype [11]. Therefore, vaginal microbiota composition might have an impact on neutrophil activation status leading to more severe CT induce immunopathology.

Microbiota composition has also an impact on CT infection. Indole produced by several anaerobic bacteria present within the vaginal microbiota, promotes CT infection in presence of IFNγ[12]. Therefore a highly diverse microbiota with a predominance of anaerobic bacteria and lack of *Lactobacillus spp*. increases the risk of CT infection[13–15]. However, a vaginal microbiota dominated by *Lactobacillus spp*. is associated with a decreased risk for CT infection[16]. An *in vitro* study demonstrated that *Lactobacillus crispatus* strains, by producing both L and D-lactic acid, have the best anti-CT effect [17]. Cynomolgus macaques, in contrast to mice, have a FRT that is closely similar to the human one in terms of morphology, endocrine system, menstrual cycle but also immune cell distribution[18,19]. In addition, macaques are susceptible to STI such as CT or Simian Immunodeficiency Virus (SIV)[20,21]. Female cynomolgus macaques have a highly diverse vaginal microbiota with few *Lactobacillus spp.* and many non-*Lactobacillus* anaerobic bacteria [22]. Their vaginal microbiota composition is thus similar to the one of dysbiotic women with a non-*Lactobacillus* dominant microbiota. The overall aim of this study was to investigate how vaginal microbiota composition influences susceptibility to *Chlamydia trachomatis* (CT) infection, as well as local innate immune responses and inflammation, including neutrophil recruitment and soluble inflammatory factors. To this end, the vaginal microbiota of female cynomolgus macaques was modified to enrich it in *Lactobacillus* species. Although this modification did not prevent CT infection, it was associated with changes in CT-specific antibody levels and inflammatory responses, indicating that vaginal microbiota composition can influence the host response to infection.

## Results

### Metronidazole treatment increases the abundance of natural *Lactobacillus spp*

To confirm vaginal microbiota alteration after antibiotic treatment and *L. crispatus* inoculations, its composition was characterized by sequencing the gene coding the 16S rRNA. At baseline, six dominant phyla were present in all animals (total mean abundance ranging from maximum to minimum): *Firmicutes* (51.58% to 92.24%), *Campylobacterota* (1.60% to 26.19%), *Proteobacteria* (0.23% to 25.83%), *Fusobacteriota* (0.78% to 20.48%), *Bacteroidota* (1.37% to 6.54%), *Actinobacteriota* (0.03% to 8.52%). The other phyla represented less than 1% of the total abundance [S1 Fig]. To evaluate the impact of metronidazole on bacterial species composition, a differentially abundant species analysis was performed comparing baseline and after antibiotic treatment. As expected, a decrease in the abundance of the main anaerobic species (*Prevotella timonensis, Peptoniphilus grossensis, Peptostreptococcus anaerobius* etc) and an increase in endogenous *Lactobacillus spp* (*L. prophage, L. johnsonii etc)* as well as some anaerobic species (*Streptococcus dysgalactiae, etc)* and aerobic species *(Staphylococcus aureus)* was observed after metronidazole treatment [Fig1; S1 Table] as described in women [23]. Then, a similar analysis was performed to study vaginal microbiota composition after two weeks of *L. crispatus* treatment, or not. *L. crispatus* was detected in the treated group only [Fig2A]. In addition, *L. crispatus* treatment induced an increase of several bacteria (*Lactobacillus reuteri, P. grossensis etc)* [Fig2A; S2 Table]. However, without *L. crispatus* treatment, animals show an increase in bacterial taxa associated with bacterial vaginosis in women such as *Peptoniphilus grossensis* and pathobiont such as *E. coli, S. aureus* as well as *Streptococcus spp* [Fig2B; S3 Table]. These results show that the effect of metronidazole is transient and species associated with human vaginosis are detected again two weeks after antibiotic treatment in untreated animals.

**Figure 1:**
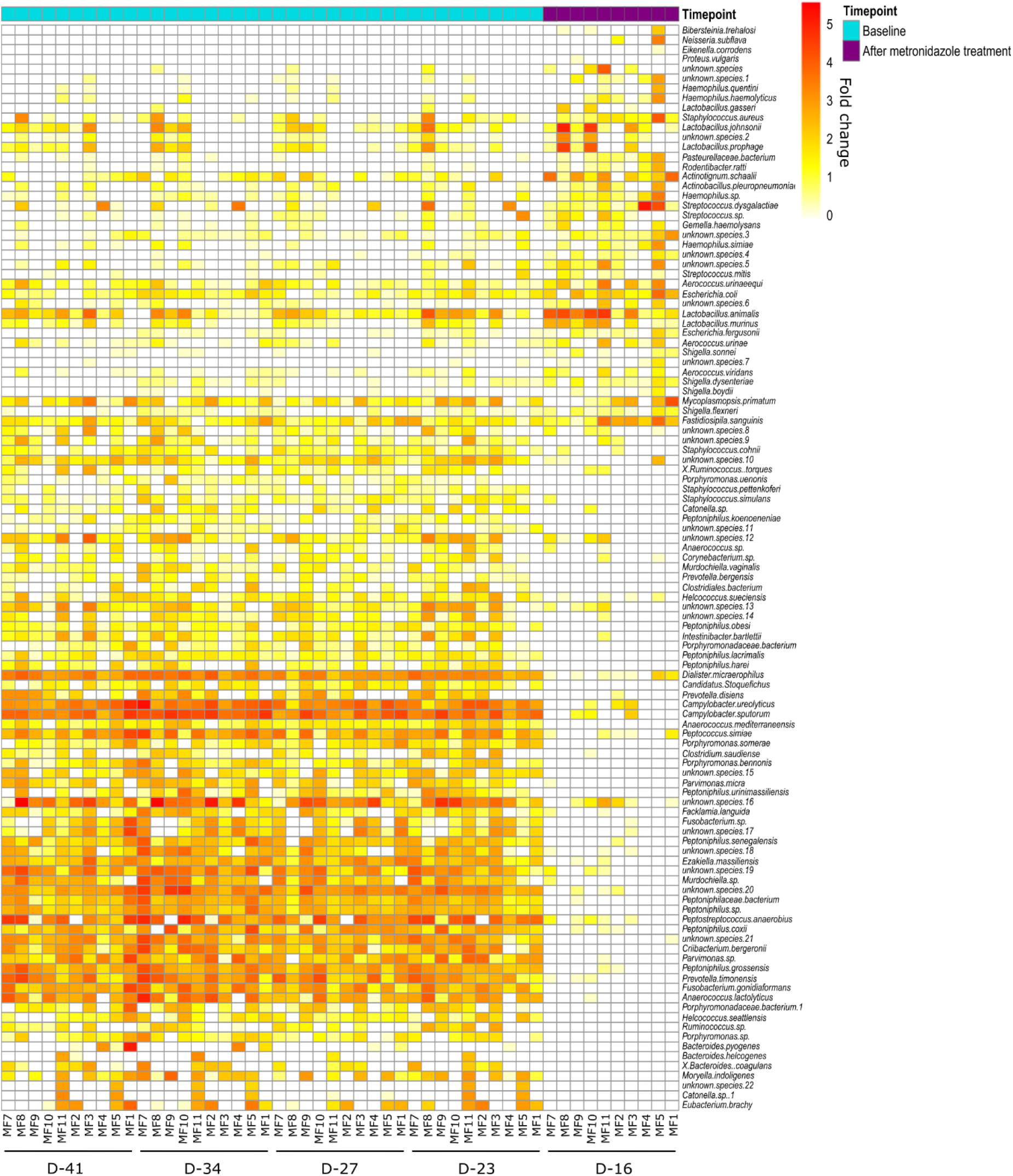
Variation of vaginal bacterial species after metronidazole treatment in all animals. DEseq2 analysis was performed to evaluate differentially absolute abundant vaginal bacterial species between the baseline and the timepoint after metronidazole treatment. Heatmap representing only the differentially abundant species is shown. The fold change after log_10_ transformation of species represents increased (from yellow to red) or decreased in the treatment condition (test condition) vs the baseline condition (control condition). Species identification are on the right of the heatmap and sample ID on the bottom. All shown species have an adjusted p-value ≤0.05.

**Figure 2:**
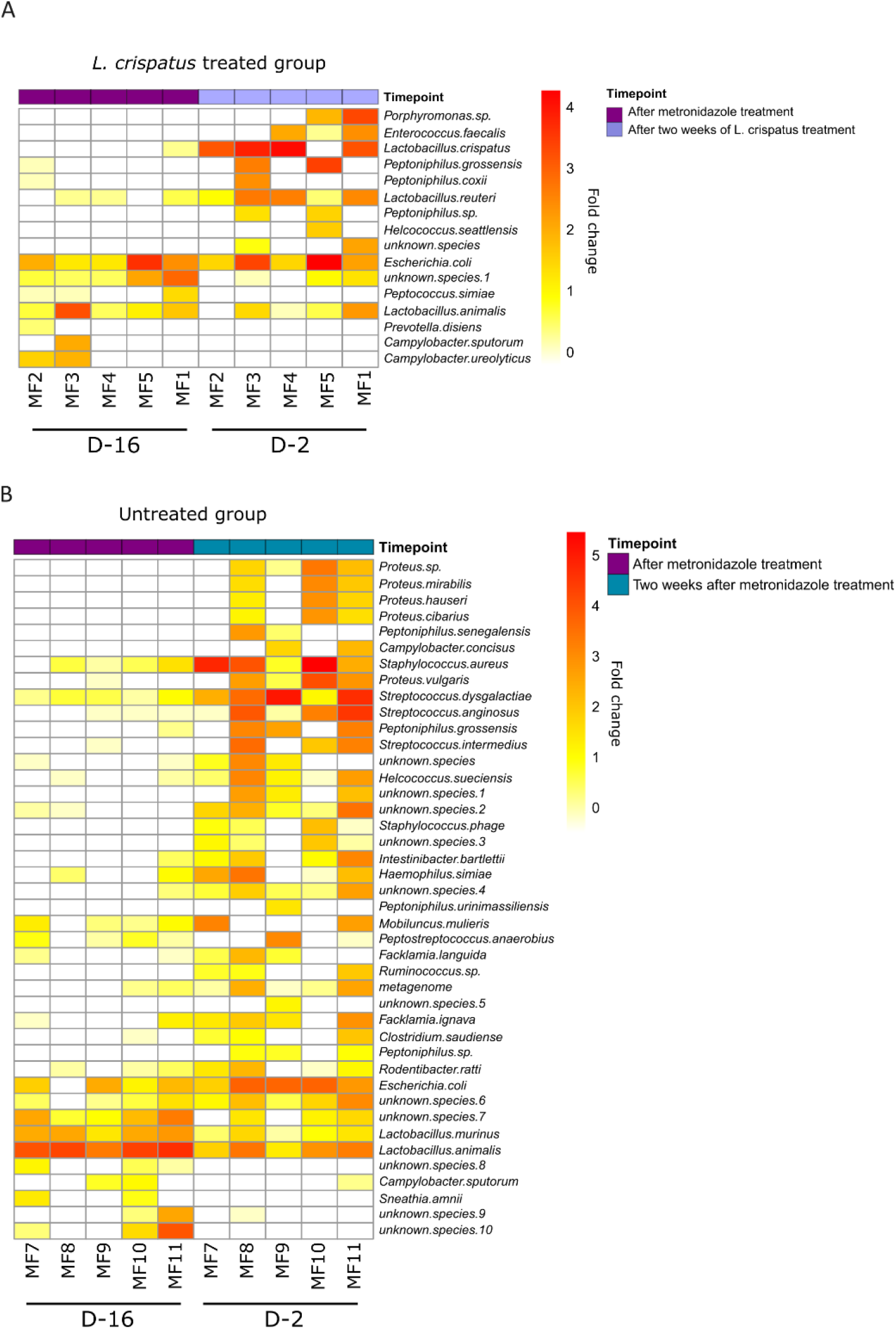
Variation of vaginal bacterial species after four inoculations of *L. crispatus* or not. DEseq2 analysis was performed to evaluate differentially abundant vaginal bacterial species between the timepoints after *L. crispatus* inoculations or not and after metronidazole treatment. Heatmap representing only the differentially absolute abundant species is shown. The fold change after log_10_ transformation of species represents increased or decreased in the treatment condition (*L. crispatus* treatment or not) vs metronidazole in the *L. crispatus* treated group (A) and in the untreated group (B). Species identification are on the right of the heatmap and sample ID on the bottom. All shown species have an adjusted p-value ≤0.05.

### Specific anti-CT IgG production is higher in *L. crispatus* treated and infected animals

To evaluate the effect of CT infection on microbiota composition, CT load quantification was performed in cervicovaginal swabs. Moreover, anti-CT immune responses were analysed by measuring serum anti-CT IgG levels and CT-specific T cells responses in PBMC after *in vitro* stimulation with inactivated CT (iCT) [Fig3A top]. Females displayed a non-persisting infection independently of their group (treated or untreated) with one or more peaks of CT load [Fig3B & C]. Area under the curve (AUC) for CT load was calculated for all individuals, and no significant differences was observed between the two groups [Fig3D, left]. Anti-CT IgGs were detected between one to three weeks after the peak of CT load [Fig3A bottom, B & C]. AUC determination for anti-CT IgG curves showed a significant higher production of antibodies in the serum of animals from the treated group [Fig3D, right]. Upon PBMC restimulation with iCT, expression of IL-2, IFNγ and TNFα by CD4+ T cells at D42 and 63 was observed without significant differences between groups [S2A & S2B Fig]. Low IL-17a expression is observed in both groups [S2B Fig]. In our experimental design, *L. crispatus* treatment once a week does not protect from CT infection. However, a higher anti-CT specific IgG response is observed in treated animals that isn’t associated with an increase in specific T cells responses in the blood [Data not shown].

**Figure 3:**
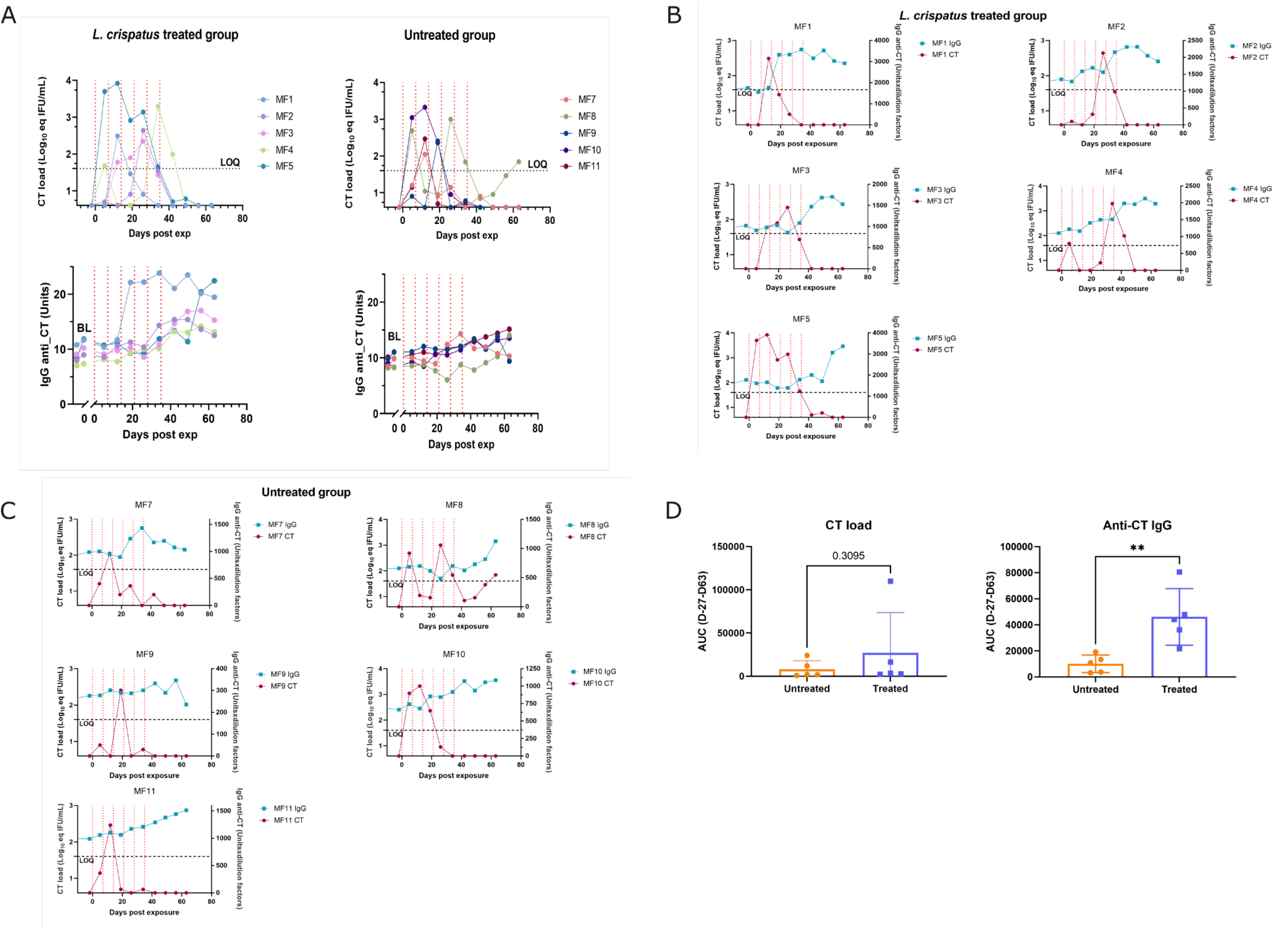
CT load and anti-CT IgG production. Graphical representation of log_10_ transformation CT load (LOD: 0.602log_10_, LOQ: 1.6log_10_) and anti-CT IgG production (Units x dilution factors) (A) in *L. crispatus* treated (left) and untreated (right) animals throughout the study. Each color represents an animal. Individual representation of CT load (log_10_ transformation) and anti-CT IgG production in each animal throughout the study in the *L. crispatus* treated group (B) and Untreated group (C). Blue and red curves match anti-CT IgG and CT load respectively. Left y axis corresponds to CT load and right y axis corresponds to anti-CT IgG. (D) Air under the curve (AUC) calculation of CT load curves (left) and anti-CT IgG (right). A Mann-Whitney test was performed to compare AUC of CT load or anti-CT IgG between groups. Asterisks indicate a p value considered statistically significant (**p ≤0.01).

### *L. johnsonii* and Lj928 abundance is associated with IgG anti-CT in *L. crispatus* untreated animals

Next, analysis of microbiota composition was performed to identify bacteria or group of bacteria that could be associated with CT load or IgG anti-CT. There was no correlation between the 9 most represented species abundances and CT load [S3 Fig]. To focus on the impact of *Lactobacillus spp* on CT susceptibility, *Lactobacillus spp* abundance, overlaid with the bacterial load was made [Fig4A and B]. *L. crispatus* abundance, observed in treated animals only, varied according to animals and time points. In the untreated group, *L. johnsonii* was the main *Lactobacillus* species observed [Fig4B]. Interestingly, peaks of CT matched with high abundance of *L. johnsonii.* A low positive association was observed between *L. johnsonii* or its prophage, Lj928, with CT load (r= 0.43 q-value: 0.0084 and r=0.45 q-value= 0.0084) [S4 Fig]. Concerning anti-CT IgG, eight and five species were positively or negatively associated with the production of anti-CT IgG, respectively [Fig4C]. Despite having a significant q-value, r coefficients are quite low. Among them*, L. johnsonii* and *Lj928* are negatively associated with antibody production (r=-0.7 and -0.73 respectively) whereas *L. crispatus* is positively associated with IgG production (r=0.37). These results demonstrated that a specific bacterial composition is involved in the regulation of anti-CT IgG levels in the sera and that *L. johnsonii* and Lj928, might be involved in high CT load and low IgG anti-CT values.

**Figure 4:**
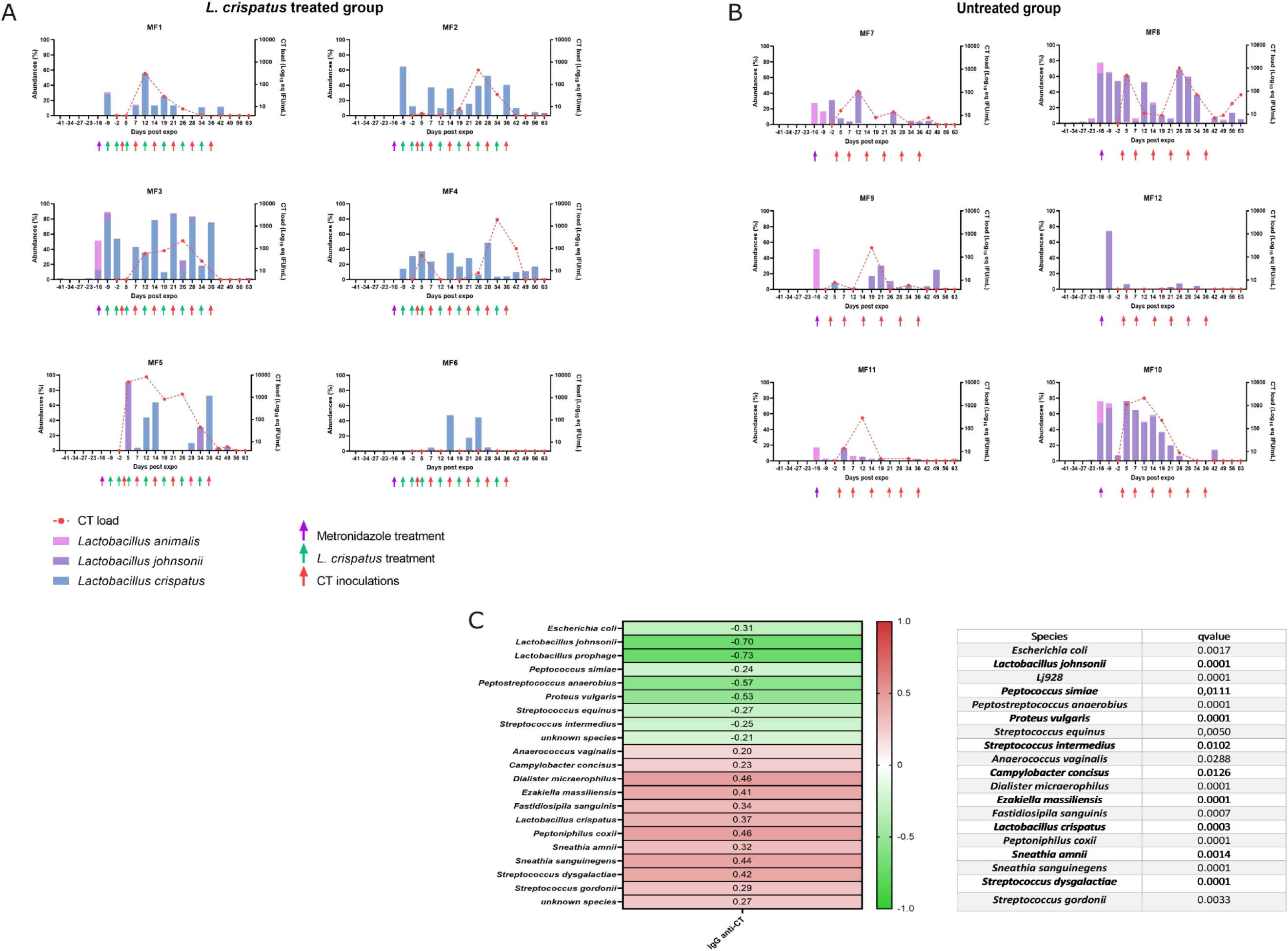
Bacterial species association with CT load and IgG anti-CT. Individual bar plot representing *Lactobacillus spp* relative abundances (%) in each animal from *the L. crispatus* treated group (A) and untreated group (B) superposed with CT load (red dotted line) after log_10_ transformation throughout the study. Purple, green and red arrows represent metronidazole treatment, *L. crispatus* inoculation and CT inoculation respectively. (C) Spearman correlation test was performed to determine association between the relative abundances of the most represented bacterial species of each animal and IgG anti-CT values. The q-values obtained are displayed on the table.

### Microbiota changes upon CT infection

Finally, to investigate bacterial taxa that were differentially abundant according to CT infection, we compared microbiota composition before, during and after CT infection. During CT infection, an increased abundance of many bacteria genus including *Anaerococcus, Blautia, Peptostreptococcus* and *Succinivibrio* is observed [Fig5A; S4 Table]. After the infection, bacterial taxa, such as *Peptoniphilus spp* and *Succinivibrio* were increased [Fig5B; S5 Table]. Thus, after metronidazole treatment there is a significant rearrangement of the vaginal microbiota even after CT infection self-resolution.

**Figure 5:**
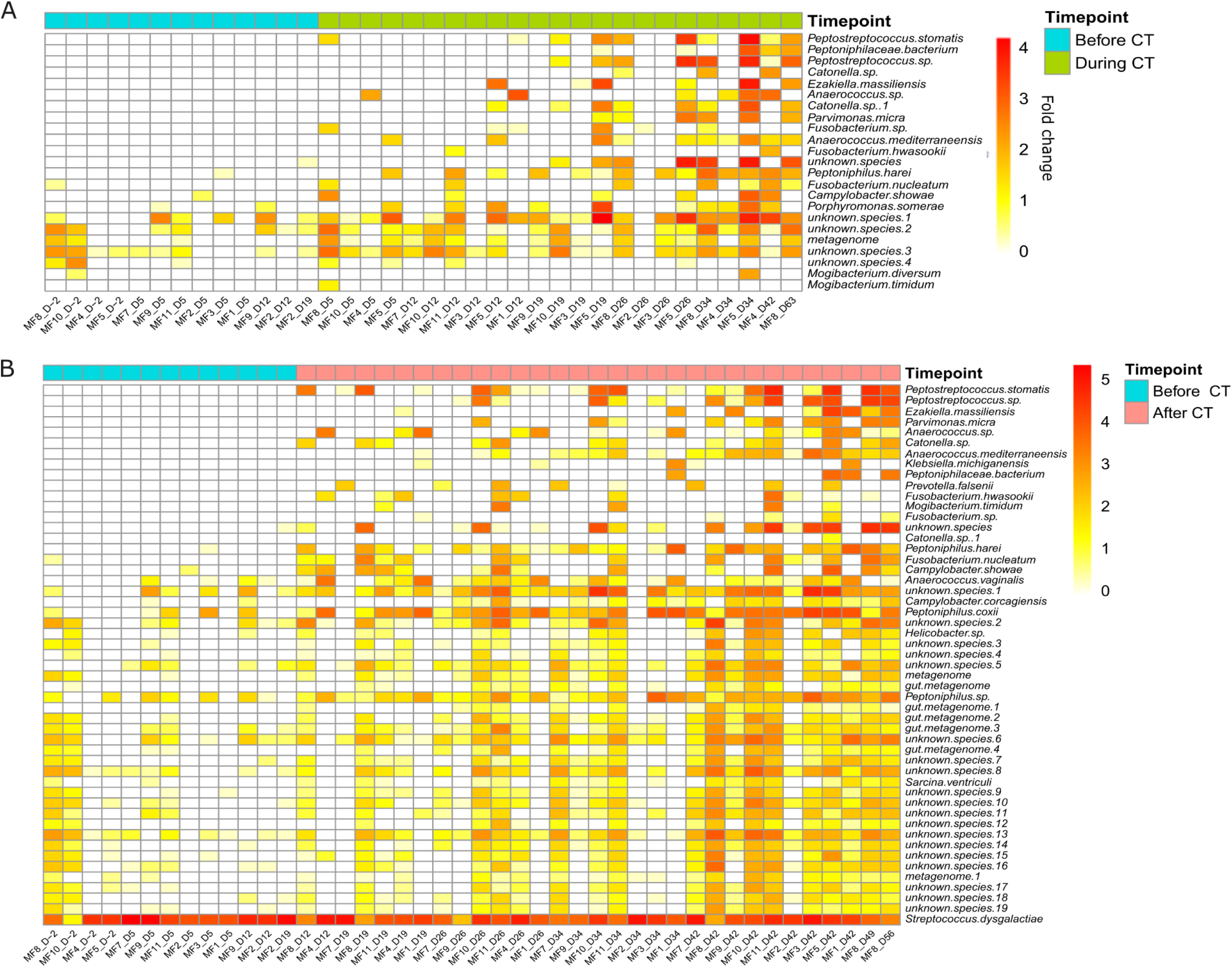
Variation of vaginal bacterial species at CT infection and after CT infection in all animals. A DEseq2 analysis was performed to evaluate differentially absolute abundant vaginal bacterial species at the infection or after CT infection compared to before CT infection (in light blue). Heatmap representing the fold change after log_10_ transformation of species increased or decreased in both conditions at during CT infection (A, in green) and after CT infection (B, in pink). The scale starts from 0 to above 4 or 5, yellow to red represents an increased abundance of the species. Species identification are on the right of the heatmap and sample ID on the bottom. All shown species have an adjusted p-value ≤0.05.

### CT infection triggered lower inflammation in the blood and higher inflammation in the cervicovaginal compartment in *L. crispatus* treated versus untreated animals

*Chlamydia trachomatis* infection induces cytokine production in the female reproductive tract [3,7] whereas a *L. crispatus* dominated microbiota is associated with a low local inflammation [24]. Therefore, to evaluate if a higher abundance of *L. crispatus* could modify CT induced cytokines/chemokines, their concentrations were measured in cervicovaginal fluids but also in the serum of all animals once a week throughout the study.

### Cytokine concentration in peripheral blood

The impact of microbiota modification on peripheral blood cytokine concentration was determined by comparing cytokine concentrations before, during and after CT infection (cf materials and methods section, S6 Table). CCL4, TGFα and IL-8 concentrations were higher in untreated animals compared to treated animal during CT infection [S5A Fig]. Moreover, most cytokines tested presented higher concentrations in untreated animals compared to treated animals after CT infection [S5B Fig]. These results demonstrate an increased cytokine production in sera after CT infection in untreated animals compare to treated animals. This higher level of cytokines after CT infection can be partially attributed to two females, MF8 and MF9 [S6 Fig].

### Cytokine expression in cervicovaginal fluids

Within cervicovaginal fluids, various cytokines including IL-1β, IL-6, IL-8 were detectable at baseline [S7 Table]. We have previously shown that the cervicovaginal environment is altered during menstruation therefore all samples collected during menstruation were excluded [25].

No significant differences were observed according to metronidazole, *L. crispatus* inoculation or CT infection in each group. However, during CT infection an increased expression of G-CSF was noticed in treated animals compared to the untreated [S5C Fig]. In addition, treated animals exhibited higher level of CCL2, G-CSF, VEGF, CCL4 and IL-15 after CT infection whereas untreated individuals had an increased expression of IL-5 compared to treated animal [S5D Fig].

Finally, no association between cytokine production in both compartments and anti-CT IgG levels were observed [Data not shown].

In human, *L. crispatus* has been associated with low local inflammation [24] but in this NHP model, during/after CT infection, higher cytokine concentrations in cervicovaginal fluids of *L. crispatus* treated compared to untreated animals was observed.

### Increase of mature and activated neutrophils after CT infection in peripheral blood but not in cervicovaginal cytobrushes

An accumulation of activated vaginal neutrophils was observed in individuals with a non-*Lactobacillus spp.* dominate microbiota [11] who have an increased risk of acquiring CT [16]. We hypothesis that activated neutrophils associated with a non-*Lactobacillus spp.* dominant microbiota could be positively associated with CT infection. Therefore, to better evaluate their role during CT infection, neutrophil phenotype was studied on whole blood and on cervicovaginal samples at five time points (two baselines, after antibiotic treatment, one week after the six CT inoculations (D42) and four weeks after the six CT inoculations (D63)). In the blood, two main populations were observed: CD11b^+^ CD101^+^ CD10^+^ CD32a^+^ and CD11b^+^ CD101^+^ CD10^-^CD32a^+^ characterized as mature and immature subsets respectively [Fig6A] as described in J. Lemaitre et al [26]. No impact of metronidazole treatment was noticed on neutrophil subpopulations [Fig6A]. In contrast, CT infection significantly increased the mature subset in both groups. Variations in the activation marker expression were evaluated on the two main neutrophil populations and compared to the post-antibiotic condition [Fig6B]. After CT infection, PD-L1 and HLA-DR are upregulated whereas CD62L and CD64 were downregulated in both groups. The immature neutrophil population showed a high variability between animals of the *L. crispatus* treated group, presenting only significant results for the downregulation of CD64 and CD62L expression [Fig6B]. Overall, no significant differences were observed between both groups of animals.

**Figure 6:**
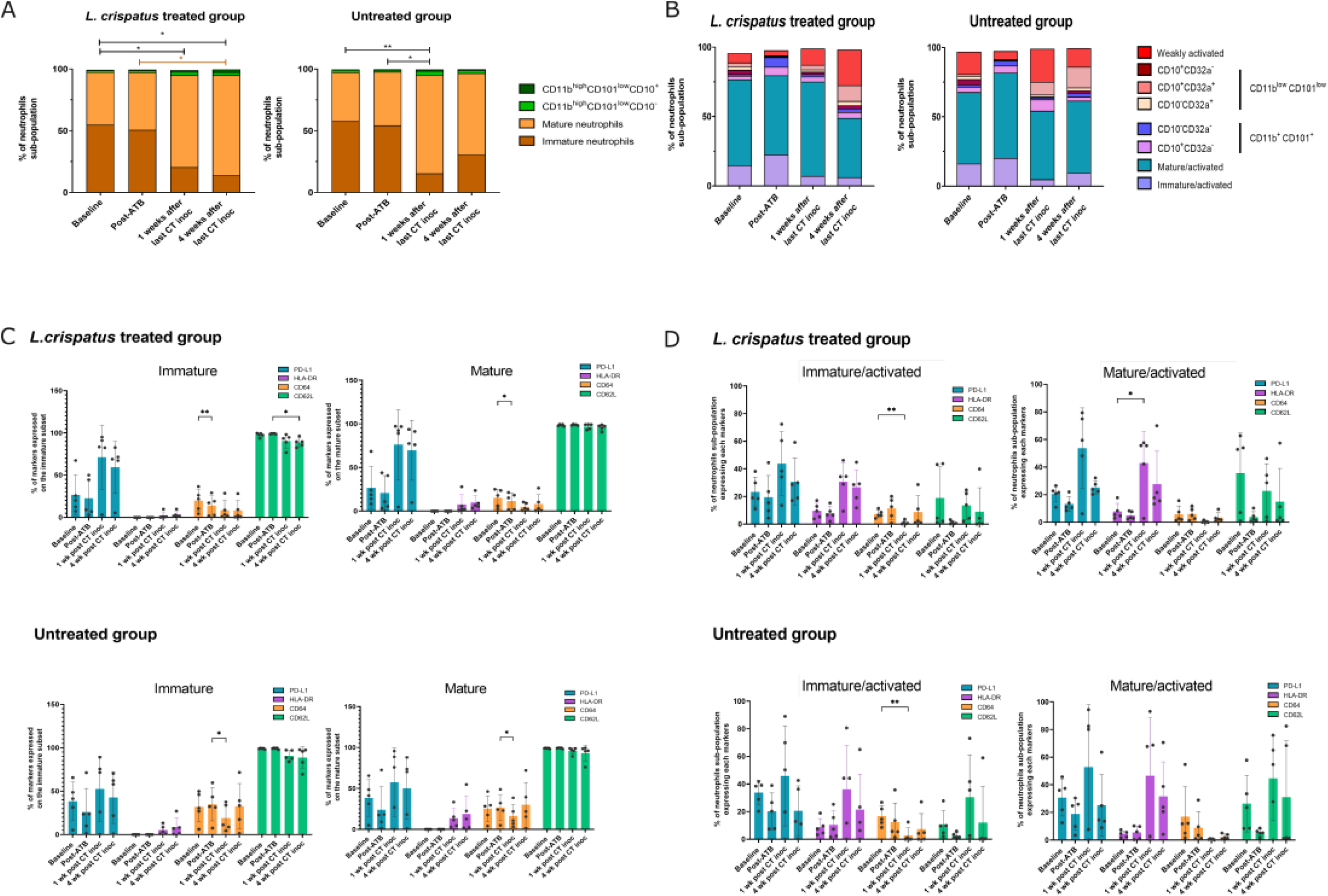
Neutrophil subpopulations and activation in peripheral blood and cervicovaginal cytobrushes. Percentage of neutrophil subpopulations among CD66+ Lin-CD14-in peripheral blood (A) and cervicovaginal cytobrushes (C) at baseline (the mean of the two baselines is represented), after metronidazole treatment or metronidazole treatment and *L. crispatus* inoculation (Post-ATB), one week after the last CT inoculation and four weeks after the last CT inoculation. Graphical representation of the percentage of the two main populations of neutrophils expressing different markers (PD-L1 in blue, HLA-DR in purple, CD64 in orange, CD62L in green) in the *L. crispatus* treated (top) or untreated (bottom) groups in the blood (B) and in the cervicovaginal compartment (C). A two-way ANOVA test with the Turkey test to adjust p value was performed to compare post-metronidazole value to other time points. Asterisks indicate a p value considered statistically significant (*p ≤0.05, **p ≤0.01).

In cervicovaginal cytobrushes, three main subsets of neutrophils were observed: mature/activated, immature/activated and weakly activated neutrophils [25] [Fig6C]. Samples collected during menstruation were excluded [25]. In terms of activation, metronidazole treatment induces a decrease of CD62L expression compared to baseline in the three main populations described above [Fig6D & S7 Fig]. After CT exposure, HLA-DR expression is significantly upregulated compared to post-antibiotic treatment (mature/activated and immature/activated population) in the treated group. Moreover, PD-L1 expression is upregulated whereas CD64 expression is downregulated. Neutrophils in both groups have a quite comparable phenotype.

In conclusion, we were unable to identify a specific neutrophil phenotype or activation status associated with microbiota composition; rather, the effects observed appear to be related to CT infection.

## Discussion

During this study, we analysed the impact of the vaginal microbiota composition on CT infection and on the cervicovaginal and peripheral blood inflammation in female cynomolgus macaques.

In our preclinical model, CT infection was detected in ten animals out of twelve and was non-persisting in contrast to what was described in pig-tailed macaques (10^6^ IFUs) [20]. The infectious dose in the present study (10^4^ IFUs) was probably not sufficient to induce a sustained CT infection. For infected animals, the T cell specific response in the blood was mainly a Th1 response with IL-2, IFNγ and/or TNFα production similar to what has been described in women able to resolve CT genital tract infection (mainly IFNγ)[4,27,28].

We showed that metronidazole treatment induced a drastic modification of the vaginal microbiota that are animal specific. Similar observations were made on Human cohort[24].

We observed that females with an *L. crispatus* enriched microbiota have a higher specific anti-CT IgG production in peripheral blood. The impact observed on IgG production was not linked to the amplitude of CT infection. In this model, various species of bacteria are either positively or negatively associated with specific anti-CT IgG production showing that a specific environment might be involved in the regulation of IgG production. In a mice model, low concentration of short chain fatty acids (SCFA) was observed to increase Ig production and Ig producing plasma cells in the gut and lymph nodes[29]. A *Lactobacillus rhamnosus* LGG derived protein was observed to promote IgA production in the gut[30]. These results demonstrated that microbiota composition through metabolite production or protein expressed on bacteria have an effect on plasma cells and Ig production. In the cynomolgus macaque model, the vaginal microbiota is similar to the one observed in dysbiotic women associated with higher SCFA concentration and lower lactic acid concentration compared to women with a *Lactobacillus spp.* dominated microbiota[22,31]. During this study, microbiota modification could have altered the proportion of SCFA or other metabolites including lactic acid, but further experiments are necessary to have a better understanding of the mechanism involving metabolites regulation of IgG production. During this study, *L. crispatus* was positively associated with IgG anti-CT whereas *L. johnsonii* and its prophage, Lj928, were negatively associated with anti-CT IgG. *Lactobacillus spp.* in women are associated with anti-inflammatory effects and protection against STI [15,16,24,32]. However, distinct inflammatory properties were observed among *Lactobacillus spp* strains, depending on whether they originated from a *Lactobacillus spp* dominant or a dysbiotic vaginal microbiota [33]. This result suggests that cynomolgus macaque *L. johnsonii* might have opposite role compared to human isolated strain of *Lactobacillus*. *L. johnsonii* abundance was not associated with a specific cytokine profile [Data not shown].

A stronger cytokine production is detected locally in treated animals compared to untreated whereas the opposite is observed in the blood. Based on the literature[24] a lower inflammation in both compartments for treated animals was expected. However, it is widely known that a balanced inflammation is needed for pathogen clearance[34] therefore we hypothesize that in this context, *L. crispatus* presence could have facilitate an efficient immune response against CT through the local production of inflammatory mediators thus decreasing the systemic inflammation. These results suggest a significant impact of vaginal microbiota composition on cytokine production locally but also in the blood as opposed to what have been observed by Hedge et al. but only few cytokines were tested in STI free individuals[35].

No differences in total neutrophil abundance was observed locally despite the increase of G-CSF in *L. crispatus* treated animals[36].

We were not able to associate a specific inflammatory profile to neutrophil recruitment or activation, locally or in the blood. Moreover, no significant differences were observed between groups in terms of neutrophil phenotype and activation. Therefore, neutrophil changes in activation/maturation status are driven by the infection and not microbiota modification. In peripheral blood, neutrophils were more mature after CT infection in both groups. The activation profile of immature/mature population is quite similar between groups associated with a non-significant increase of PD-L1 and HLA-DR and significant decrease of CD64 post-infection. During CT infection, neutrophils were described to have a delay in apoptosis[10,37] which could be mediated through the increased expression of PD-L1 as observed in a mouse model of sepsis[38]. In cervicovaginal cytobrushes, CD62L expression on the three main neutrophil populations was decreased after metronidazole treatment as observed on colonic neutrophils of mice treated with an antibiotic cocktail[39]. However, no effect of antibiotic treatment was detected on cytokine production (systemic and local) or T cell specific response on PBMC. CD62L downregulation after antibiotic treatment can be associated to an increased activation, also observed after infection[39,40]. No impact on maturation was noticed after CT exposure in contrast to peripheral blood neutrophils possibly due to a late sampling, far from CT infection. Neutrophils were more activated and have a prolonged lifespan as observed in the blood.

This is the first study describing neutrophils in blood and cervicovaginal cytobrushes in a context of CT infection.

Our study has some limitations. Significant inter-individual differences enable us to evaluate the effect of vaginal microbiota on CT infection, therefore a higher number of animals should be investigated for further studies. We were able to reproduce an asymptomatic CT infection observed in women but a more severe model with a sustained infection would be interested to study. Future investigations aiming to clarify the respective contributions of CT infection and antibiotic exposure on vaginal microbiota alterations may consider including a CT-infected, metronidazole-untreated group.

Overall, this study demonstrates that female macaques with a *L. crispatus* enriched vaginal microbiota developed a stronger specific antibody response against CT in blood compared to *L. crispatus* untreated animal. Moreover, a specific microbiota composition, after metronidazole treatment, induce a weaker specific immune responses associated with a lower anti-CT IgG concentration. Interestingly, systemic cytokine levels were higher in untreated animals, while treated animals showed increased cytokines production in the cervicovaginal compartment, indicating a more effective local immune response.

In conclusion, this study demonstrates a significant impact of the vaginal microbiota composition on the local and systemic immune response induced by CT infection.

## Materials and methods

### Ethics approval

Twelve sexually mature adult female cynomolgus macaques (*Macaca fascicularis*), aged of 5 to 7 years old and originating from Mauritian AAALAC certified breeding centers were included in this study. All animals were housed in IDMIT facilities (CEA, Fontenay-aux-roses), under BSL-2 containment (Animal facility authorization #D92-032-02, Préfecture des Hauts de Seine, France) and in compliance with European Directive 2010/63/EU, the French regulations and the Standards for Human Care and Use of Laboratory Animals, of the Office for Laboratory Animal Welfare (OLAW, assurance number #A5826-01, US). Animals undergo a 40-day acclimatization before study entry. This study was approved and accredited by the institutional ethical committee “Comité d’Ethique en Expérimentation Animale du Commissariat à l’Energie Atomique et aux Energies Alternatives” (CEtEA #44) under statement numbers A18-083. The study was authorized by the “Research, Innovation and Education Ministry” under registration number APAFIS#20692-2019051709424034v1. The twelve animals were housed in the same room in social groups of six animals under controlled conditions of humidity, temperature and light (12h light/dark cycles). The animals were fed twice a day with commercial monkey chow and fruits. Water was available *ad libitum*. They were provided with environmental enrichment including toys and novel food under the supervision of the CEA Animal Welfare Officer.

### Experimental design

Twelve female cynomolgus macaques (MF1 to MF12) divided in two groups (Untreated and *L. crispatus* treated) were studied. Age, weight, haplotype and group randomization are summarized in S1 Table. Randomization was done using excel. According to the data previously obtained by our team [S8 Fig], the following protocol was selected to modify the microbiota composition. *L. crispatus* strain isolated from a commercially available compound (Physioflor^®^, IPRAD SANTÉ, Gentilly, France), observed to decrease by 20.5% the recurrence rate of BV [41], was used during this study. The experimental design is represented in the Fig7A. On days -23, -21 and -19, all females received a reconditioned vaginal ovule of metronidazole (FLAGYL 500mg; SANOFI, Paris, France). Then, a *L. crispatus* gel was inoculated intravaginally to the females of the treated group. All the animals received intravaginally 10^4^ IFU of *C. trachomatis* serovar D once a week during six weeks. Then animals were followed up for four weeks [Fig7A]. After CT inoculations, two animals (MF6 from the *L. cripatus* treated group and MF12 from the untreated group) did not get infected nor developed a specific anti-CT immune response (serum anti-CT IgG or T cell). Moreover, no specific vaginal microbiota composition could be associated with the absence of CT infection. Therefore, the results of these two animals are not shown in the manuscript.

### Sample collection

Whole blood, cervicovaginal fluids and swabs were collected [Fig7A] at baseline, after metronidazole treatments, after one and two weeks of *L. crispatus* inoculations or not, two days before each CT inoculation and once a week during the follow up. All time points after metronidazole treatments will be referred as “post-antibiotic treatment”; *L. crispatus* treated animals will be referred as “treated” and *L. crispatus* untreated animals as “untreated”. Samples collection are specified in Fig7A. Sample collection order was as followed: Weck-Cel Spear during blood withdrawal, vaginal pH measurement, cervicovaginal swab and cervicovaginal cytobrushes. Samples included: cervicovaginal fluids to measure cytokine/chemokine concentration; blood used for Complete Blood Count (CBC), cytokine/chemokine and progesterone quantification; cervicovaginal samples for Nugent score determination and microbiota characterization and cervicovaginal cytobrushes for neutrophil phenotyping were collected as previously described[25]. Cell Preparation tube (CPT) with Sodium Heparin were used to isolate mononuclear cells from peripheral blood (BD Biosciences, Le Pont-de-Claix, France).

Samples collected during the experiment (from D0 to D63):

• Before CT infection will be referred as “before CT”

• During CT infection (CT load >1.6 log_10_ eq IFU/mL) will be referred as “during CT”

• After CT infection (after CT infection & when the CT load <1.6 log_10_ eq IFU/mL) will be referred as “after CT”.

### Metronidazole ovules

Metronidazole ovules (Flagyl^®^, Sanofi, Paris, France) of 500mg active substance were put in a 5 mL tube and weight before putting in a water bath set at 37°C until complete melting. Melted ovules were then used to fill pediatric ovule adapted to the size of the macaque vaginal cavity (Herboristerie du Valmont; Belgium). Ovules were then weighted to determine the concentration of each antibiotic ovule (187.7 mg ±17.8/ovule (mean±SD))) and stored at 4°C before inoculation.

### L. crispatus gel

*L. crispatus* (Physioflor LP^®^; IPRAD SANTÉ, Paris, France) strain (IP174178) was cultured in Mann Rogosa and Sharpe (MRS) liquid culture media (BD Bioscience, Le Pont-de-Claix, France) supplemented with 0.05% L-cysteine (Sigma Aldrich) overnight at 37 °C with 10 % CO_2_. Inoculums were harvested as previously described [42]. Briefly, bacteria were collected 1h after reaching the stationary phase, washed twice with PBS 1X and resuspended in 1.5% Hydroxyethylcellulose (HEC) gel (Sigma Aldrich, Missouri, Saint-Louis, USA), 9% sucrose (Sigma Aldrich), 0.05% lactic acid (Sigma Aldrich) and citrate buffer (Citric acid monohydrate; trisodium citrate dihydrate; H_2_O milliQ). Syringes were filled with 1 mL of *L. crispatus* gel and stored at -80 °C until use. Animals were inoculated with 10^10^ CFUs. Serial dilutions of the inoculum were plated in MRSA to verify the inoculum after each thawing.

### C. trachomatis infection

*Chlamydia trachomatis* serovar D (D/UW3/Cx) were obtained from Dr Frank Follmann laboratory (SSI, Danemark) [43]. For animal inoculation, CT was diluted in Sucrose Phosphate Glutamate (SPG) buffer to achieve the expected concentration. Each animal received intravaginally 800µL of CT inoculum (10^4^ IFU/800 µL).

### Progesterone quantification

Progesterone level was determined in plasma sample by ELISA (IBL international; Hambourg, Germany) according to manufacturer’s instructions. Animals showed normal menstrual cycles, with most exhibiting ≥3 progesterone peaks; only two (MF1 and MF5) had fewer [Fig 7B].

**Figure 7:**
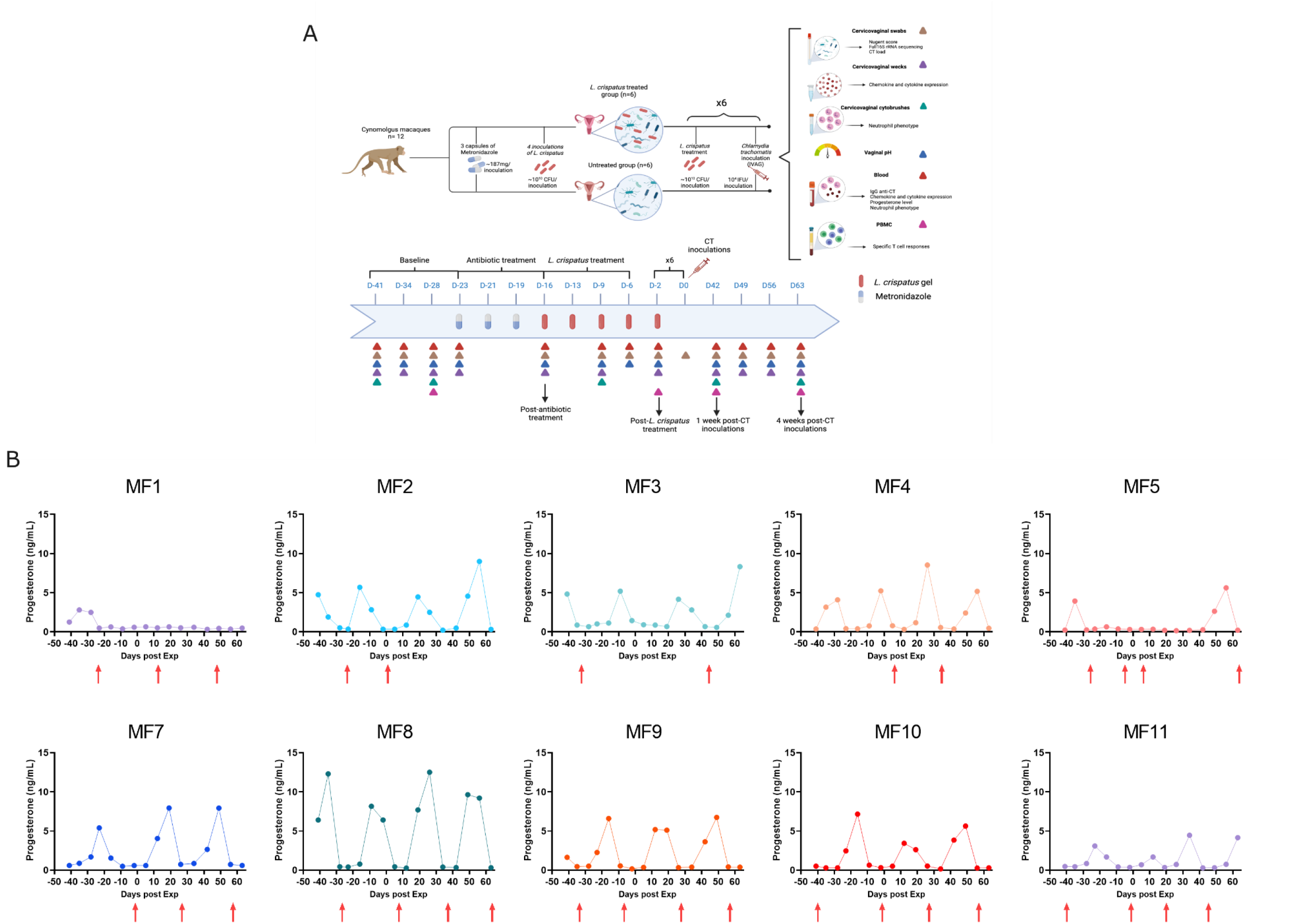
Experimental design and progesterone level. (A) Twelve female cynomolgus macaques were included in this study and divided in two groups of six animals. After four weeks of baseline, all animals received intravaginally three ovules of metronidazole during one week. Then animals from the *L. crispatus* treated group received four inoculations of *L. crispatus* gel at 10^10^ CFU (twice during two weeks) after washing the vaginal cavity with 0.02% of lactic acid. At D0, all animals were inoculated with 10^4^ IFU of CT. At D-2 all animals within the *L. crispatus* treated group received the gel of *L. crispatus*. This design D-2/D0 was repeated six times, therefore all animals received six doses of CT. Then animals were studied during four weeks post-infection. Samples collection included: cervicovaginal swabs, cervicovaginal wecks, cervicovaginal cytobrushes, whole blood, PBMC. The pH was also measured. Created with Biorender.com. (B) Individual plots of the progesterone quantification (ng/mL) throughout the study are shown for the 10 infected animals. Red arrows represent menstruation.

### Chlamydia trachomatis load

Detection of CT in cervicovaginal samples was performed by qPCR. DNA was extracted using the kit QIAamp DNA Mini Kit (Qiagen, Hilden, Germany) according to manufacturer’s instructions. As for qPCR, the kit Presto combined quantitative real time *Chlamydia trachomatis/Neisseria gonorrhea* (Goffin Molecular Technologies, Waalwijk, Netherlands) and a thermocycler CFX96 Touch Real-Time PCR detection system (Biorad, California, Hercules, USA) was used according to manufacturer’s instructions. Limit of detection (LOD) is fixed at 0.602 log_10_ and limit of quantification (LOQ) at 1.6 log_10_.

### Specific antibodies against CT

Specific IgG against CT were quantified by ELISA (Demeditec; Kiel, Germany) in serum according to manufacturer’s instructions. Goat anti-monkey IgG coupled with HRP at 50 ng/mL (Biorad, California, Hercules, USA) was used for revealing. Optical density values at 450nm were obtained using a multimode microplate reader, the Spark 10M (TECAN, Männedorf, Switzerland). Units of IgG anti-CT was calculated using the following formula: (OD_sample_ x10) / OD_cutoff_ and multiplied by the dilution factor.

### DNA extraction and full 16S rRNA gene sequencing

The PowerFecal DNA Pro isolation kit (Qiagen, Hilden, Germany) was used following the manufacturer’s instructions. 16S rRNA gene was amplified by PCR using specific 16S primers (27F: 5’ AGAGTTTGATCMTGGCTCAG-3′ and 1492R: 5′- CGGTTACCTTGTTACGACTT-3′) of the 16S Barcoding Kit 1-24 (SQK-16S024, Oxford Nanopore Technologies) and LongAmp™ Taq 2× Master Mix (New England Biolabs, Evry-Courcouronnes, France). PCR conditions used were 1 min of denaturation at 95 °C, 30 cycles of denaturation (20 secs at 95 °C), annealing (30 secs at 55 °C) and extension (2 min at 65 °C). A final extension step was added for 5 min at 65 °C. PCR products were washed using AMPure XP beads (Beckman Coulter, Indianapolis, Indiana, USA) and quantified using QuBit fluorometer and HS DNA kit (Invitrogen). Twenty-four barcoded samples were pooled and 100 fmol (∼100 ng) were used for library preparation on one R9.4 flow cell. GridION ™ was used for sequencing (Oxford Nanopore Technologies, Oxford, Great Britain) according to the manufacturer’s instructions.

### Cytokine and chemokine quantification

Pro- and anti-inflammatory cytokines as well as chemokines were quantified in cervicovaginal fluids and plasma as previously described [25].

### Neutrophil phenotyping

Neutrophil populations were analyzed in whole blood and cells obtained from cervicovaginal cytobrushes as previously described [25]. A fourteen-color panel, containing neutrophil surface makers of maturation and activation, were used (S9 Table). The gating strategy used is described in the S9 Fig.

### Peripheral T cell responses

PBMC were plated with co-stimulation markers (CD28 and CD49d) and Brefeldin A (10µg/mL; Sigma-Aldrich) in addition to different stimuli overnight at 37°C: unstimulated, PMA (6,2µg/mL; Sigma-Aldrich) and ionomycin (72µg/mL; Sigma-Aldrich) or inactivated *Chlamydia trachomatis* (iCT; 5µg/mL) (ProteoGenix, Schiltigheim, France). Cells were stained with a viability dye (Live/Dead Fixable Blue dead cell stain kit; ThermoFisher), then permeabilized using Cytofix/Cytoperm reagent (BD Biosciences) and Perm/Wash buffer (BD Biosciences) and freezed for each timepoint. Antibody staining was performed in a single step following storage and permeabilization. The antibodies used for extra- and intra-cellular staining are listed in the S10 Table and a LSRII with DIVA (BD Biosciences) and FlowJo software (Tristar) were used for the analysis.

### Microbiota and Statistical analysis

Sequence reads were converted into FASTQ files. A quality filter using NanoFilt was applied. Reads were filtered on quality metrics (Q > 10) and on read length (reads between 1200-1800 bp were kept. SILVA138_16S_pintail_100 (<u>Index of/frogs_databanks/assignation/SILVA/16S (inra.fr)</u>) was used for assignation using Minimap2 (2.17-r954-dirty) for alignment. A supplementary filter was applied on alignment metrics and on the percentage of identity. Species abundance was obtained per sample (reads assigned to one species/total reads of the sample). Raw abundance file was converted to a biom file and Phyloseq object was created using FROGS pipeline (Find Rapidly OTU with Galaxy Solution) [44] implemented on a galaxy instance (http://migale.jouy.inra.fr/galaxy/). Finally, to study differentially abundant species in different conditions, we used DESeq2 [45]. Heatmaps representing transformed fold change of cytokine/chemokine concentration in cervicovaginal fluids or plasma were obtained by Tableau software (Seattle, USA). GraphPad prism software version 9 for Windows (GraphPad Software, La Jolla California USA, www.graphpad.com) was used for graphical representation of the vaginal microbiota, cytokine concentration, and neutrophil subpopulations. Significant differences between groups were confirmed using either a Mann Whitney, a two-way ANOVA test with p value adjustment with Turkey test or Kruskal-Wallis test with p values adjustment with Dunn’s test.

## Acknowledgements

The program was funded by the Infectious Disease Models and Innovative Therapies (IDMIT) research infrastructure supported by the “Programme Investissements d’Avenir”, managed by the ANR (ANR-11-INBS-0008), as well as Sidaction “financement jeune chercheur” (2019-2-FJC-1234). The Non-Human Primate study is part of the TracVac project, which received financial support from the European Union’s Horizon 2020 research and innovation programme under grant agreement No. 733373.

The authors would like to thank all members of the ASW (Raphaël Ho Tsong Fang; Jean-Marie Robert, Sebastien Langlois, Maxime Potier, Quentin Sconosciuti, Nina Dhooge), L2I (Julie Morin) and LFC (Jerôme Van Wassenhove, Mario Gomez-Pacheco) teams of the IDMIT infrastructure as well as Animalliance group members (Thierry Prot, Marjorie Benfissa). The authors thank Dr Agathe Subtil (Institut Pasteur, department of cellular biology of microbial infection, Paris) for the discussion surrounding this project, Dr Luc de Chaisemartin (Institut Pasteur, Inserm UMR1222, APHP Bichat hospital) for his help on neutrophil data interpretation and Dr Frank Follmann as well as his team in Statens Serum Institut (Copenhagen, Denmark) for their assistance and provision of the CT strain used in this study.

## Supporting information captions

**S1 Fig. Vaginal microbiota composition at the phylum level.** Percentage of mean relative abundances of phylum is represented in pie chart for each animal *L. crispatus* treated animals (left) and untreated animals (right) at baseline.

**S2 Fig. CD4⁺ T-Cell Polyfunctionality.** (A) Graphical representation of the percentage of CD4+ T cells expressing Th1 cytokines (IL-2, TNFα, IFNγ) at baseline, D42 and D63 with iCT *in vitro* stimulation or not (NS) in *L. crispatus* treated (blue) and untreated animal (orange). (B) Graphical representation of the percentage of CD4+ T cells expressing Th1 cytokines (IL-2, TNFα, IFNγ) or IL-17A at baseline, D42 and D63 with iCT *in vitro* stimulation in *L.crispatus* treated (blue) and untreated group (orange).

**S3 Fig. Vaginal microbiota composition at the species level throughout the study**. Percentage of relative abundances of the top 9 most represented species, in *L. crispatus* treated animals (left) and untreated animals (right), is represented in bar plot. The red dotted line represents CT load in each animal.

**S4 Fig. Association between *L. johnsonii* or *L. johnsonii prophage* (*Lj928*) relative abundance and CT load.** (A) Heatmap representing the Spearman correlation coefficient was generated. Green represents negative association and red a positive association. The q values obtained are displayed on the table. (B) Simple linear regression was performed to compare the relative abundance of *L. johnsonii* or *Lj928* with CT load. Goodness of fit is shown with r^2^.

**S5 Fig. Cytokine quantification in peripheral blood and cervicovaginal fluids.** Log_10_ transformation of cytokine concentrations in peripheral blood (A and B) and in cervicovaginal fluids (C and D) of untreated animals (orange) and *L. crispatus* treated (blue) animals during CT infection (A and C) or after CT infection (B and D). A Mann- Whitney test was performed to compare cytokine production between groups. Asterisks indicate a p-value considered statistically significant (*p ≤0.05, **p ≤0.01, ***p ≤0.001, **** p ≤0.0001).

**S6 Fig. Peripheral blood cytokine concentrations in each animal.** Heatmap representing the fold change of cytokines and chemokines in peripheral blood serum in each animal throughout the study. The fold change was calculated based on the expression of each cytokine/chemokine for each female at baselines. Red indicates an increased fold change and blue a decrease.

**S7 Fig. Expression of Activation Markers on Weakly Activated Cervicovaginal Neutrophils.** Graphical representation of the percentage of weakly activated cervicovaginal neutrophils expressing different markers (PD-L1 in blue, HLA-DR in purple, CD64 in orange, CD62L in green) in the *L. crispatus* treated (top) or untreated (bottom) groups.

**S8 Fig. Strategies tested to enrich the vaginal microbiota with *L. crispatus*.** Several protocols were tested to induce the enrichment of exogenous *Lactobacillus spp*. in five female cynomolgus macaques : (A) A single inoculation of a capsule of *L. crispatus* (Physioflor LP^®^; IPRAD PHARMA, Paris) ; (B) A single pretreatment with metronidazole two days before the inoculation of *L. crispatus* capsule; (C) Depo-provera® intra-muscular injection (30mg) one month before metronidazole pretreatment two days before the inoculation of *L. crispatus* capsule (Physioflor LP^®^; IPRAD PHARMA, Paris); (D) Depo-provera® intra-muscular injection (30mg) in 3 animals out of 6, one month before repeated sucrose gel inoculations; (E) Repeated treatments of metronidazole before repeated inoculations of *L. crispatus* capsule (Physioflor LP^®^; IPRAD PHARMA, Paris); (F) Repeated treatments of azithromycine before repeated inoculations of *L. crispatus* gel. Menstruation and Depo-provera injections reduced the enrichment of *L. crispatus*. However, repeated pretreatments with metronidazole or azithromycin followed by repeated inoculations of *L. crispatus* induced a higher rate of *L. crispatus* enrichment. Created with Biorender.com.

**S9 Fig. FACS gating strategy for neutrophil phenotyping**. (A) In the peripheral blood samples and (B) cervicovaginal cytobrushes.

**S1 Table. Affiliation of unknown species at the genus or family level associated with the Fig1.**

**S2 Table. Affiliation of unknown species at the genus or family level associated with the Fig2A.**

**S3 Table. Affiliation of unknown species at the genus or family level associated with the Fig2B.**

**S4 Table. Affiliation of unknown species at the genus or family level associated with the Fig5A.**

**S5 Table. Affiliation of unknown species at the genus or family level associated with the Fig5B.**

**S6 Table. Cytokine concentrations (pg/ml) at baseline (n=1) in the peripheral blood in each animal.**

**S7 Table. Median cytokine concentrations (pg/ml) at baseline (n=4) in cervicovaginal fluids in each animal.**

**S8 Table. Information concerning the 12 animals included in the study.**

**S9 Table. Antibody panel for neutrophil phenotyping in blood and cervicovaginal cytobrushes.**

**S10 Table. Antibody panel for T cell intracellular staining.**

## DATA AVAILABILITY

rRNA16S sequences associated with this project are deposited in SRA under BioProject accession number PRJNA1152928. Other datasets generated during and/or analyzes during the current study are available from the corresponding author on reasonable request.

## AUTHORS’ CONTRIBUTIONS

The study was conceived and designed by CA, LR, NN, RM, JL, RLG, MTN, and EM. Data acquisition was performed by CA, LR, CC, ML, LB, WG, and MTN. Data analysis and interpretation was done by CA, LR, NN, SL, SD, LDA, WG, ASG, ML, LB, NDB, MTN, and EM. CA and EM drafted the manuscript. Critical revisions was performed by CA, LR, NN, RM, JL, RLG, MTN, ASG and EM.

