## Supplementary material for "The vaginal microbiota shapes local and systemic inflammation during *Chlamydia trachomatis* infection in non-human primate": S1 Table

| ***Species*** | ***Genus*** | ***Family*** |
|---|---|---|
| Unknown species | *Catonella* | |
| Unknown species 1 | *Peptoniphilus* | |
| Unknown species 2 | *Anaerococcus* | |
| Unknown species 3 | *Parvimonas* | |
| Unknown species 4 | *Fusobacterium* | |
| Unknown species 5 | *Shuttleworthia* | |
| Unknown species 6 | *Peptostreptococcus* | |
| Unknown species 7 | *Facklamia* | |
| Unknown species 8 | *Terrisporobacter* | |
| Unknown species 9 | *Sarcina* | |
| Unknown species 10 | *Clostridium sensu stricto 1* | |
| Unknown species 11 | *Porphyromonas* | |
| Unknown species 12 | *Fastidiosipila* | |
| Unknown species 13 | *Negativibacillus* | |
| Unknown species 14 | *Helcococcus* | |
| Unknown species 15 | *Eschirichia-Shigella* | |
| Unknown species 16 | *Aerococcus* | |
| Unknown species 17 | *Actinobacillus* | |
| Unknown species 18 | *Unknown genus* | *Actinomycetaceae* |
| Unknown species 19 | *Staphylococcus* | |
| Unknown species 20 | *Aggregatibacter* | |
| Unknown species 21 | *Lactobacillus* | |
| Unknown species 22 | *Haemophilus* | |
| Unknown species 23 | *Anaerobiospirillum* | |
