## Supplementary material for "The vaginal microbiota shapes local and systemic inflammation during *Chlamydia trachomatis* infection in non-human primate": S2 Table

| ***L. crispatus* treated group** | | |
|---|---|---|
| **Species** | **Genus** | **Family** |
| Unknown species | *Shuttleworthia* | |
| Unknown species 1 | *Succinivibrio* | |
| Unknown species 2 | Unknown genus | *Actinomycetaceae* |
| Unknown species 3 | *Sarcina* | |
| Unknown species 4 | *Gemella* | |
