## Supplementary material for "The vaginal microbiota shapes local and systemic inflammation during *Chlamydia trachomatis* infection in non-human primate": S3 Table

| Untreated group | |
|---|---|
| **Species** | **Genus** |
| Unknown species | *Anaerobiospirillum* |
| Unknown species 1 | *Parvimonas* |
| Unknown species 2 | *Ruminococcus* |
| Unknown species 3 | *Anaerococcus* |
| Unknown species 4 | *Facklamia* |
| Unknown species 5 | *Terrisporobacter* |
| Unknown species 6 | *Staphylococcus* |
| Unknown species 7 | *Helcococcus* |
| Unknown species 8 | *Peptoniphilus* |
