## Supplementary material for "The vaginal microbiota shapes local and systemic inflammation during *Chlamydia trachomatis* infection in non-human primate": S4 Table

| Before CT vs during CT | |
|---|---|
| **Species** | **Genus** |
| Unknown species | *Aggregatibacter* |
| Unknown species 1 | *Blautia* |
| Unknown species 2 | *Succinivibrio* |
| Unknown species 3 | *Anaerococcus* |
| Unknown species 4 | *Parvimonas* |
