## Supplementary material for "The vaginal microbiota shapes local and systemic inflammation during *Chlamydia trachomatis* infection in non-human primate": S5 Table

| Before CT vs after CT | | |
|---|---|---|
| **Species** | **Genus** | **Order** |
| Unknown species | *Christensenellaceae R7 group* | |
| Unknown species 1 | *UCG-002* | |
| Unknown species 2 | *Marvinbryantia* | |
| Unknown species 3 | *Rikenellaceae RC9 gut group* | |
| Unknown species 4 | *Peptococcus* | |
| Unknown species 5 | *[Ruminococcus] gauvreauii group* | |
| Unknown species 6 | *UCG-008* | |
| Unknown species 7 | *Oribacterium* | |
| Unknown species 8 | *Terrisporobacter* | |
| Unknown species 9 | *Subdoligranulum* | |
| Unknown species 10 | *Dorea* | |
| Unknown species 11 | *Blautia* | |
| Unknown species 12 | *Unknown genus* | *Clostridia UCG-014* |
| Unknown species 13 | *Sarcina* | |
| Unknown species 14 | *Clostridum senso stricto 1* | |
| Unknown species 15 | *Fournierella* | |
| Unknown species 16 | *Prevotella_9* | |
| Unknown species 17 | *Succinivibrio* | |
| Unknown species 18 | *Anaerococcus* | |
| Unknown species 19 | *Parvimonas* | |
