## Supplementary material for "The vaginal microbiota shapes local and systemic inflammation during *Chlamydia trachomatis* infection in non-human primate": S6 Table

| | G-CSF | IFNγ | IL-1RA | IL-2 | IL-5 | IL-6 | IL-8 | IL-10 | IL-12/23(p40) | IL-13 | IL-15 | CCL2 | CCL3 | CCL4 | TGFα | TNFα | VEGF |
|---|---|---|---|---|---|---|---|---|---|---|---|---|---|---|---|---|---|
| **MF1** | 50.04 | 1.93* | 16.08 | 20.93 | 2.25* | 9.58* | 6002.89 | 12.21* | 26.04 | 9.73* | 9.72* | 213.41 | 2.51* | 12.06 | 4.65 | 2.44* | 16.9* |
| **MF2** | 25.73 | 1.93* | 24.4 | 11.22 | 2.25* | 9.58* | 1412.77 | 12.21* | 10.13* | 9.73* | 9.72* | 237.29 | 2.51* | 19.06 | 3.92 | 2.44* | 16.9* |
| **MF3** | 11.5* | 4.42 | 36.2 | 19.98 | 2.25* | 9.58* | 682.9 | 12.21* | 10.13* | 9.73* | 9.72* | 513.15 | 2.51* | 26.56 | 7.56 | 2.44* | 91.17 |
| **MF4** | 11.5 | 1.93* | 44.61 | 16.47 | 2.25* | 9.58* | 477.76 | 12.21* | 10.13* | 9.73* | 9.72* | 373.38 | 2.51* | 14.36 | 8.91 | 2.44* | 16.9* |
| **MF5** | 29.7 | 9.95 | 453.47 | 21.85 | 2.25* | 9.58* | 846.45 | 111.03 | 10.13* | 9.73* | 9.72* | 757.15 | 9.7 | 15.45 | 2.46 | 2.44* | 38.29 |
| **MF7** | 42.79 | 4.62 | 17.95 | 15.3 | 2.56* | 2.42* | 5069.47 | 13.59* | 2.73* | 9.8* | 8.48 | 390.56 | 6.94 | 21.49 | 12.45 | 2.44* | 45.06 |
| **MF8** | 26.62 | 9 | 28.24 | 44.38 | 7.15 | 2.91 | 6096.75 | 22.8 | 2.73* | 9.8* | 7.62 | 888.44 | 6.94 | 19.93 | 12.34 | 14.46 | 151.39 |
| **MF9** | 34.97 | 9 | 29.08 | 28.67 | 2.56* | 3.84 | 4491.47 | 25.91 | 52.04 | 12.05 | 6.16 | 560.98 | 10.85 | 22.88 | 3.33 | 53.54 | 106.06 |
| **MF10** | 26.62 | 2.59* | 20.31 | 16.09 | 2.56* | 2.42* | 1633.58 | 13.59* | 2.73* | 9.8* | 2.58* | 99.29 | 2.42* | 29.43 | 19.65 | 2.44* | 24.15 |
| **MF11** | 29.47 | 9 | 54.13 | 32.09 | 2.56* | 2.42* | 3166.21 | 13.59* | 6.89 | 9.8* | 7.62 | 582.7 | 5.33 | 25.23 | 5.88 | 262.39 | 70.46 |

*Out of range values
