## Supplementary material for "The vaginal microbiota shapes local and systemic inflammation during *Chlamydia trachomatis* infection in non-human primate": S7 Table

| | G-CSF | IL-1RA | IL-1β | IL-2 | IL-5 | IL-6 | IL-8 | IL-10 | IL-12/23 (p40) | IL-13 | IL-15 | IL-18 | CCL2 | CCL3 | CCL4 | sCD40L | TGFα | TNFα | VEGF |
|---|---|---|---|---|---|---|---|---|---|---|---|---|---|---|---|---|---|---|---|
| **MF1** | 205.645 | 4077.505 | 9.68 | 2.38 | 2.38 | 235.24 | 2529.155 | 11.01 | 2.03 | 4.325 | 3.18 | 39.16 | 209.195 | 3.835 | 49.21 | 12.46 | 15.83 | 36.73 | 66.25 |
| **MF2** | 99.58 | 4704.57 | 21.63 | 2.36 | 2.38 | 4.2 | 2219.07 | 11.66 | 1.88 | 2.875 | 3.695 | 34.85 | 229.915 | 3.685 | 13.17 | 29.36 | 7.83 | 26.71 | 113.985 |
| **MF3** | 25.48 | 3482.47 | 6.595 | 2.36 | 2.38 | 2.36 | 789.52 | 11.66 | 1.88 | 2.32 | 2.31 | 23.675 | 18.015 | 2.37 | 4.105 | 3.76 | 7.235 | 10.76 | 4.97 |
| **MF4** | 222.34 | 10202.65 | 143.915 | 2.38 | 2.38 | 79.645 | 1253.52 | 21.14 | 5.445 | 5.36 | 7.19 | 242.285 | 263.6 | 4.81 | 38.495 | 12.46 | 12.08 | 56.125 | 312.29 |
| **MF5** | 59.355 | 7072.19 | 10.285 | 2.38 | 2.38 | 21.835 | 2084.57 | 24.69 | 2.03 | 3.965 | 4.4 | 21.88 | 259.945 | 3.195 | 29.79 | 49.415 | 19.79 | 77.925 | 85.44 |
| **MF7** | 257.895 | 9658.31 | 33.905 | 2.32 | 2.44 | 6.115 | 2033.99 | 13.49 | 4.35 | 3.78 | 2.795 | 438.365 | 82.1 | 2.99 | 15.075 | 36.94 | 6.185 | 10.025 | 59.78 |
| **MF8** | 74.655 | 9658.31 | 6.15 | 3.64 | 17.88 | 6.07 | 3453.39 | 23.79 | 5.36 | 5.075 | 3.755 | 72.935 | 93.3 | 6.155 | 6.525 | 137.685 | 8.11 | 14.72 | 98.635 |
| **MF9** | 301.31 | 9658.31 | 57.24 | 2.375 | 2.44 | 37.3 | 2764.425 | 13.49 | 3.33 | 4.1 | 4.195 | 94.095 | 240.73 | 2.4 | 27.6 | 28.05 | 6.005 | 17.81 | 121.905 |
| **MF10** | 29.7 | 9658.31 | 4.75 | 2.32 | 2.44 | 2.495 | 2400.54 | 13.49 | 2.31 | 2.48 | 2.62 | 79.95 | 131.775 | 2.4 | 4.06 | 4.2 | 7.76 | 9.37 | 28.855 |
| **MF11** | 57.395 | 4265.28 | 17.41 | 2.36 | 2.38 | 3.365 | 1423.215 | 11.66 | 1.88 | 2.32 | 2.31 | 29.515 | 31.505 | 2.37 | 7.235 | 3.76 | 2.875 | 10.76 | 14.51 |
