## Supplementary material for "The vaginal microbiota shapes local and systemic inflammation during *Chlamydia trachomatis* infection in non-human primate": S8 Table

| ID | Date of birth | Weight (Kg) | Haplotype | Groups |
|---|---|---|---|---|
| MF1 | 2015-02-05 | 3.59 ± 0.09 | Rec H4 H7<br>Rec H4 H7 | *L. crispatus* treated group |
| MF2 | 2015-03-24 | 5.47 ± 0.27 | H3<br>H2 | *L. crispatus* treated group |
| MF3 | 2015-04-12 | 4.97 ± 0.27 | Rec H3 H4 H7<br>Rec H7 H1 H2 | *L. crispatus* treated group |
| MF4 | 2015-04-19 | 4.21 ± 0.09 | H1<br>Rec H1 H2 | *L. crispatus* treated group |
| MF5 | 2015-08-08 | 4.22 ± 0.25 | H1<br>H2 | *L. crispatus* treated group |
| MF6 | 2015-08-27 | 4.78 ± 0.33 | H1<br>H6 | *L. crispatus* treated group |
| MF7 | 2015-02-06 | 2.84 ± 0.06 | Rec H4 H7<br>Rec H3 H1 H2 | Untreated group |
| MF8 | 2015-03-22 | 5.31 ± 0.27 | H1<br>Rec H4 H5 | Untreated group |
| MF9 | 2015-04-06 | 4.64 ± 0.33 | H1<br>Rec H2 H4 H7 | Untreated group |
| MF10 | 2015-08-06 | 4.70 ± 0.17 | Rec H1 H4 H7<br>H3 | Untreated group |
| MF11 | 2015-08-11 | 6.75 ± 0.31 | H3<br>Rec H1 H5 | Untreated group |
| MF12 | 2015-08-18 | 3.35 ± 0.08 | Rec H3 H4<br>Rec H7 H1 H3 H6 | Untreated group |
