## Supplementary material for "The vaginal microbiota shapes local and systemic inflammation during *Chlamydia trachomatis* infection in non-human primate": S9 Table

| Antibody (clone) | Label | Reference | Manufacturer |
|---|---|---|---|
| Bluevid | BUV736 | L23105 | Life Technologies |
| CD64 (10.1) | BUV737 | 612776 | BD |
| CD11b (REA713) | FITC | 130-110-552 | Miltenyi |
| CD45 (REA1023) | Viogreen | 130-177-193 | Miltenyi |
| CD3 (SP34.2 | BV650 | 563916 | BD |
| CD8 (RPAT8) | BV650 | 563821 | BD |
| CD20 (2H7) | BV650 | 563780 | BD |
| CD123 (7G3) | BV650 | 563405 | BD |
| CD62L (SK11) | BV711 | 565040 | BD |
| CD14 (REA599) | Vioblue | 130-110-524 | Miltenyi |
| CD10 (HI10a) | PercP-Cy5.5 | 312216 | BioLegend |
| CDw125 (REA705) | PE | 130-110-544 | Miltenyi |
| PD-L1 (29E.2A3) | PE-Dazzle594 | 329732 | BioLegend |
| CD101 (REA954) | PE-Vio770 | 130-115-832 | Miltenyi |
| CD32a (IV.3) | AF647 | 60012 | Stem Cell |
| HLA-DR (L234) | AF700 | 307626 | BioLegend |
| CD66 (TET2) | APC-Vio770 | 130-119-847 | Miltenyi |
