## Supplementary material for "The vaginal microbiota shapes local and systemic inflammation during *Chlamydia trachomatis* infection in non-human primate": S10 Table

| Antibody | Label | Reference | Manufacturer |
|---|---|---|---|
| Bluevid | BUV736 | L23105 | Life Technologies |
| IFNγ | V450 | 560371 | BD |
| CD4 | BV510 | 563094 | BD |
| TNFα | BV605 | 502936 | Bio Legend |
| IL-13 | BV711 | 564288 | BD |
| CD154 | FITC | 555699 | BD |
| IL-5 | PE | 559332 | BD |
| IL-2 | PerCP-Cy5.5 | 560708 | BD |
| CD8 | PE-Vio770 | 130-113-159 | Miltenyi |
| CD137 | APC | 550890 | BD |
| IL-17A | AF700 | 560613 | BD |
| CD3 | APC-Cy7 | 557757 | BD |
